# The conserved molting/circadian rhythm regulator NHR-23/NR1F1 serves as an essential co-regulator of *C. elegans* spermatogenesis

**DOI:** 10.1101/2020.06.11.147298

**Authors:** James Matthew Ragle, Abigail L. Aita, Kayleigh N. Morrison, Raquel Martinez-Mendez, Hannah N. Saeger, Guinevere A. Ashley, Londen C. Johnson, Katherine A. Schubert, Diane C. Shakes, Jordan D. Ward

## Abstract

In sexually reproducing metazoans, spermatogenesis is the process by which uncommitted germ cells give rise to haploid sperm. Work in model systems has revealed mechanisms controlling commitment to the sperm fate, but how this fate is subsequently executed remains less clear. While studying the well-established role of the conserved nuclear hormone receptor transcription factor, NHR-23/NR1F1, in regulation of *C. elegans* molting, we discovered NHR-23/NR1F1 is also constitutively expressed in developing 1° spermatocytes and is a critical regulator of spermatogenesis. In this novel role, NHR-23/NR1F1 functions downstream of the canonical sex determination pathway. Degron-mediated depletion of NHR-23/NR1F1 within hermaphrodite or male germlines causes sterility due to an absence of functional sperm as depleted animals produce arrested primary spermatocytes rather than haploid sperm. These spermatocytes arrest in prometaphase I and fail to either progress to anaphase or attempt spermatid-residual body partitioning. They make sperm-specific membranous organelles (MOs) but fail to assemble their major sperm protein into fibrous bodies. NHR-23/NR1F1 appears to function independently of the known SPE-44 gene regulatory network, revealing the existence of an NHR-23/NR1F1-mediated module that regulates the spermatogenesis program.

**Summary Statement:** A well-characterized regulator of *C. elegans* molting also unexpectedly controls the spermatogenesis program; our work provides insights into the gene regulatory networks controlling spermatogenesis.

## Introduction

Transcription factors perform diverse roles in the development of specialized cell types. Some function as switches to direct binary decisions about cell fate (Davis et al., 1987; Gualdi et al., 1996; Horner et al., 1998; Isshiki et al., 2001). Others work synergistically in gene regulatory networks to execute complex programs of cell differentiation (Davidson and Levin, 2005; Medwig-Kinney et al., 2020; Olson, 2006; Peter and Davidson, 2011). These networks coordinate the expression of enzymes, modifiers, and structural proteins that characterize a specific cell type. One challenge of studying transcription factors is they often function with distinct co-regulators in multiple developmental contexts. For example in the nematode *Caenorhabditis elegans*, the GATA-1 transcription factor ELT-1 is essential for epidermal cell fates during embryogenesis (Page et al., 1997) but later regulates the expression of multiple sperm genes (del Castillo-Olivares et al., 2009). In knockout or null mutants, if disruption of the first developmental event results in embryonic or larval lethality, later functions will be missed. Traditionally this challenge has been addressed using conditional mutants, mRNA depletion by RNA interference, or genetic mosaics. A powerful new addition to this investigational toolbox is the auxin-inducible degron (AID) system for the conditional disruption of protein function (Nishimura et al., 2009; Zhang et al., 2015). In this system, a gene of interest tagged with an AID sequence is co-expressed with a TIR1 F-box protein driven by a tissue-specific promoter. In the presence of the plant hormone, auxin, TIR1 polyubiquitinates the AID sequence, leading to degradation of the AID-tagged protein. This conditional protein depletion system is ideal for characterizing the distinct functions of an essential, multi-functional transcription factor.

One motivation of developing the AID system in *C. elegans* was to study events in meiosis required for gametogenesis (Zhang et al., 2015). During gametogenesis, stem cell precursors enact a developmental program producing highly specialized haploid sperm or oocytes. *C. elegans* is a powerful model to study gametogenesis as hermaphrodites produce a limited number of sperm before switching exclusively to producing oocytes whereas males produce sperm continuously (Ellis and Schedl, 2007)(Fig. 1A,B). Extensive studies of *C. elegans* sex determination (Barton and Kimble, 1990; Ellis and Schedl, 2007) identified the transcription factor TRA-1, a homolog of GLI and cubitus interruptus, as the key regulator of somatic sex determination and the spermatocyte/oocyte decision (Hodgkin, 1987; Schedl et al., 1989; Zarkower and Hodgkin, 1992). Within the germline, TRA-1 promotes oogenesis/inhibits spermatogenesis by repressing expression of two germline-specific, RNA-binding proteins (FOG-1 and FOG-3)(Chen and Ellis, 2000; Jin et al., 2001). A different RNA-binding translational repressor, PUF-8, maintains sperm fate (Subramaniam and Seydoux, 2003).

**Fig. 1.**
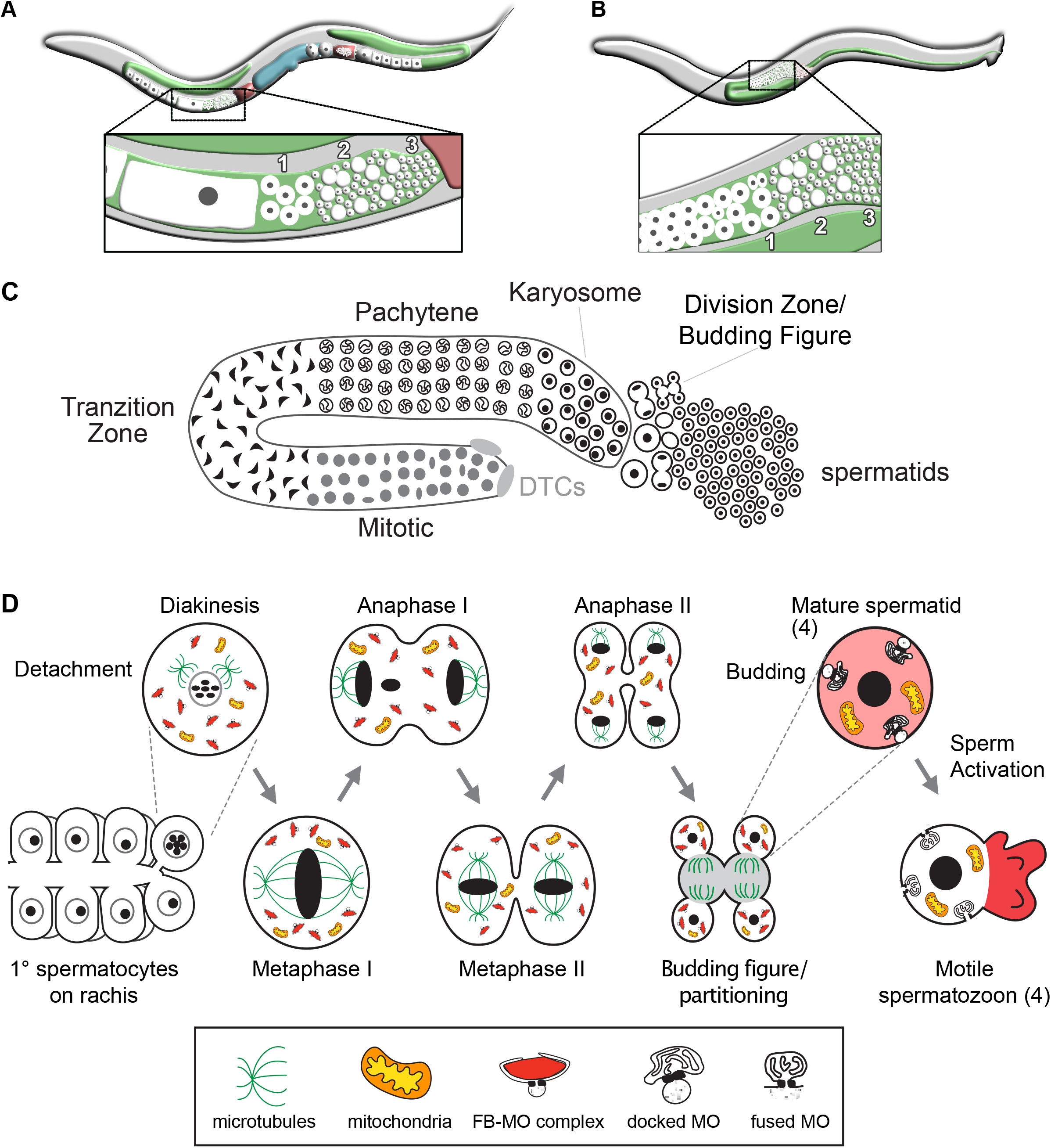
Overview of *C. elegans* spermatogenesis. (A,B) Cartoons depicting a young adult *C. elegans* hermaphrodite and male and their respective germlines. The hermaphrodite germline (A) is transitioning from spermatogenesis to oogenesis. The enlarged views highlight the linear arrangement of the primary spermatocytes (1), residual bodies (RB) (2), and mature haploid spermatids (3). (C) Stylized cartoon of a surface view of the male germline highlighting its overall linear organization. Mitotic proliferation of the germline stem cells is maintained by two somatic distal tip cells (DTCs) which form the germ cell niche. The early events of meiotic prophase including homolog pairing and formation of the synaptonemal complex, occur in the transition zone. Following an extended pachytene stage, spermatocytes enter a karyosome stage before mature spermatocytes detach from the syncytial germline and divide meiotically. The first meiotic division is often incomplete, leaving secondary spermatocytes linked by a cytoplasmic connection. Following anaphase II, the spermatocytes morph into budding figures that split into residual bodies and haploid spermatids. (D) Details of the meiotic divisions and post-meiotic partitioning event. Once spermatocytes detach from the germline syncytium, they pass through a brief diakinesis stage before undergoing nuclear envelope breakdown and initiating meiotic divisions. During the post-meiotic partitioning event, microtubules become acentrosomal and localize to the developing residual body (Winter et al., 2017). Components retained in the spermatids include fibrous body-membranous organelles (FB-MO), mitochondria, chromatin, and centrioles. Components discarded within the RB include the cell’s tubulin, actin, ER, and ribosomes; mature sperm are thus both transcriptionally and translationally inactive. Following separation from the RB, FBs disassemble to release their MSP and the MOs dock with the plasma membrane. Males store sperm in this inactive spermatid state. During sperm activation, MOs fuse with the plasma membrane and MSP localizes to the pseudopod.

In contrast to our understanding of how sperm fate is controlled, the control of sperm differentiation is much less clear. Understanding the gene regulatory networks that control sperm differentiation offers an inroad into addressing this question. The *C. elegans* transcription factor SPE-44 is widely distributed on autosomes of developing spermatocytes and is required for proper expression of one-quarter of the spermatogenesis transcriptome (Kulkarni et al., 2012). *spe-44* directly regulates other transcription factors like *elt-1* as well as sperm function genes such as kinases and structural components (Kulkarni et al., 2012). Which other transcription factors contribute to gene expression regulation in *C. elegans* spermatogenesis and to what degree they control similar or disparate sets of genes remains unknown.

Spermatogenesis is a complex cellular process involving a host of dynamic subcellular events coordinated by a large number of genes. Because of its linear organization, the full developmental sequence of spermatogenesis can be analyzed in individual gonads (Fig. 1C, Chu and Shakes, 2013). Undifferentiated germ cells proliferate mitotically at the distal end while mature primary spermatocytes at the proximal end divide meiotically before undergoing the post-meiotic event of spermatid-residual body separation (Figure 1C,D). The commitment to sperm fate occurs as undifferentiated germ cells exit mitosis and enter the initial stages of meiotic prophase within the transition zone. Subsequent to homolog pairing, transcription of all spermatogenesis genes and translation of most sperm proteins occurs within an extended pachytene zone. Near the end of meiotic prophase, global transcription ceases as developing spermatocytes enter a karyosome stage during which chromosomes detach from the nuclear envelope and coalesce into a central mass (Shakes et al., 2009). Although transcription stops, translation and assembly of sperm-specific complexes continues (Chu and Shakes, 2013; Shakes et al., 2009). Throughout meiotic prophase, individual spermatocytes exist in a syncytium, linked to a shared central rachis (Fig. 1D). Following the karyosome stage, primary spermatocytes detach from this rachis before undergoing the meiotic divisions. Post-meiotic development is limited to a partitioning event that occurs as spermatids bud from a central residual body and a sperm activation process during which spherical spermatids polarize and acquire motility through extension of a pseudopod (Chu and Shakes, 2013; Ward et al., 1981). Despite having large collections of mutants affecting the processes of sperm differentiation (Nishimura and L’Hernault, 2010), how these events are coordinated is not well-understood.

*C. elegans* NHR-23/NR1F1 (hereafter referred to as NHR-23) is a nuclear hormone receptor (NHR) transcription factor previously shown to be involved in embryonic epidermal development (Kostrouchova et al., 1998), larval molting (Kostrouchova et al., 1998; Kostrouchova et al., 2001), and diet-induced developmental acceleration (Macneil et al., 2013). The mammalian orthologs (RORs/NR1F) regulate circadian rhythms (Akashi and Takumi, 2005; André et al., 1998; Sato et al., 2004), and the insect ortholog (DHR3) regulates metamorphosis (Kageyama et al., 1997; Lam et al., 1997; White et al., 1997). Loss or reduction of *nhr-23* activity leads to embryonic lethality, larval arrest, or larval lethality (Kostrouchova et al., 1998; Kostrouchova et al., 2001; Kouns et al., 2011). Lethality following *nhr-23* inactivation occurs early in development, masking an unanticipated later role for NHR-23 in spermatogenesis which we describe here. Our results are the first to show that *nhr-23* is expressed within developing spermatocytes and functions downstream of the canonical sex determination pathway. Depletion of NHR-23 protein results in sperm-specific sterility. Affected spermatocytes arrest in meiosis I prometaphase and exhibit developmental defects in biogenesis of an essential sperm-specific organelle. NHR-23 promotes spermatogenesis in a distinct pathway from SPE-44. Our work provides new insights into how transcription factors coordinately execute the spermatogenesis program.

## Results

### NHR-23:GFP is expressed in sperm-producing germlines

To explore potential functions of NHR-23 beyond molting, we wanted to know where and when NHR-23 is expressed. To address this, we used CRISPR/Cas9-mediated genome editing to introduce a multifunctional GFP::AID*::3xFLAG sequence in the endogenous *nhr-23* locus, tagging all known isoforms. AID* is a minimal, 44 amino acid auxin-inducible degron sequence, and the 3XFLAG epitope is useful for western blotting. Consistent with previous promoter reporters and immunolabelling (Kostrouchova et al., 1998), NHR-23::GFP expression was observed in epidermal cell nuclei of 1.5-stage embryos and in hypodermal and seam cells of developing larvae (unpublished data). Somatic cell expression persisted in L4 larvae, specifically in hypodermal cells of the head, vulval precursor cells of hermaphrodites, and hypodermal and tail cells of males (Fig. 2A). Notably, NHR-23::GFP was also expressed in the sperm-producing germlines of males and L4 hermaphrodites, but not in hermaphrodites only producing oocytes (Fig. 2B). Previous studies missed this spermatocyte expression as transgenes are frequently silenced in the germline and immunolabelling was limited to embryos (Kostrouchova et al., 1998; Kostrouchova et al., 2001).

**Fig. 2.**
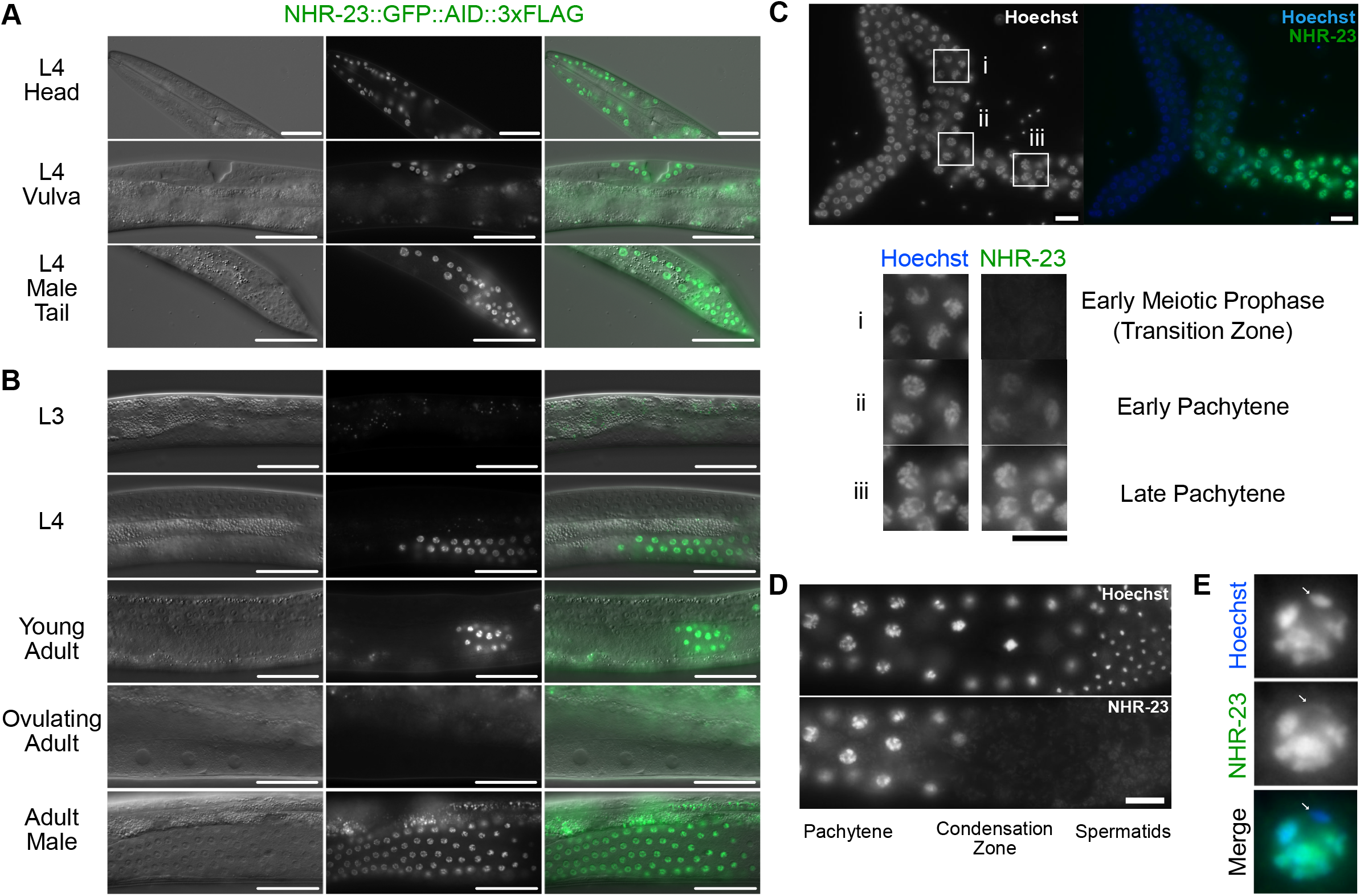
NHR-23::GFP::AID*::3xFLAG is expressed in somatic cells throughout development and in spermatogenic germlines. A strain carrying GFP::AID*::3xFLAG knocked-in to the endogenous *nhr-23* gene to produce a C-terminal translational fusion to all known *nhr-23* isoforms was used to monitor endogenous NHR-23 expression. (A) Representative DIC and GFP images of NHR-23::GFP::AID*::3xFLAG L4 larvae, specifically hypodermal cells of a hermaphrodite head, vulval precursor cells, and seam/hypodermal cells of the male tail. (B) Representative DIC and GFP images of NHR-23::GFP::AID*::3xFLAG in pachytene cells of the germline in L3, L4, young adult, ovulating adult and adult male worms. (C) Fluorescent images of NHR-23::GFP::AID*::3xFLAG in a dissected adult male germline. Inset boxes show cells in early meiotic prophase (transition zone) (i), early (ii) and late (iii) pachytene. (D) Representative fluorescent images of NHR-23::GFP::AID*::3xFLAG in a late pachytene male germline. (E) Representative image of a late pachytene nucleus expressing NHR-23::GFP::AID*::3xFLAG. The location of the X chromosome is noted with a white arrow. Scale bars: (A, B) 40 μm, (C-E) 10 μm. A minimum of 12 P0 animals were used in A, B, and C. Nuclei were visualized by Hoechst stain in C-E.

In males (and sperm-producing hermaphrodites), NHR-23::GFP was first observed in early pachytene spermatocytes (Fig. 2C), increased in intensity through late pachytene (Fig. 2B, C), and became undetectable during late meiotic prophase with onset of chromatin condensation (Fig. 2D). NHR-23 was undetectable in meiotically dividing spermatocytes or mature spermatids. This overall pattern is similar to that reported for SPE-44 (Kulkarni et al., 2012). Also like SPE-44, NHR-23::GFP labeled most chromosomes along their entire length yet failed to label a portion of DNA in each pachytene stage spermatocyte (Fig. 2E). Presumably NHR-23 is not labelling the X chromosome, which is transcriptionally silent during spermatocyte meiosis (Kelly et al., 2002).

### Auxin-induced NHR-23-depletion during or before spermatogenesis causes infertility in males and hermaphrodites

Because both *nhr-23* null mutants and RNAi knockdowns of L1 larvae arrest as early larvae (Kostrouchova et al., 1998; Kostrouchova et al., 2001), previous studies failed to assess potential NHR-23 functions in the germline. Conversely, RNAi treatment of L3/L4 hermaphrodites or adult males failed to detectably deplete NHR-23 in the germline (unpublished data). We therefore turned to auxin-inducible degradation (Nishimura et al., 2009; Zhang et al., 2015), which allows for tissue-specific, temporally regulated depletion of proteins. Our multifunctional NHR-23 knock-in construct contained the AID* sequence necessary for auxin- and TIR-dependent depletion of target proteins. A separate TIR1 transgene included the strong *mex-5* promoter (Schubert et al., 2000) which would allow NHR-23-depletion primarily in the germline. Hermaphrodites expressing either *mex-5::TIR1* or *nhr-23::GFP::AID*::3xFLAG* grew to adulthood and were fully fertile whether grown on control or 4 mM auxin media (Fig. 3A). Hermaphrodites with both constructs were fully fertile on control media but produced no progeny when grown on media with 4 mM auxin. Crossing NHR-23-depleted hermaphrodites to wild type males restored fertility (Fig. 3B). These results suggest NHR-23 is necessary for sperm production in hermaphrodites.

**Fig. 3.**
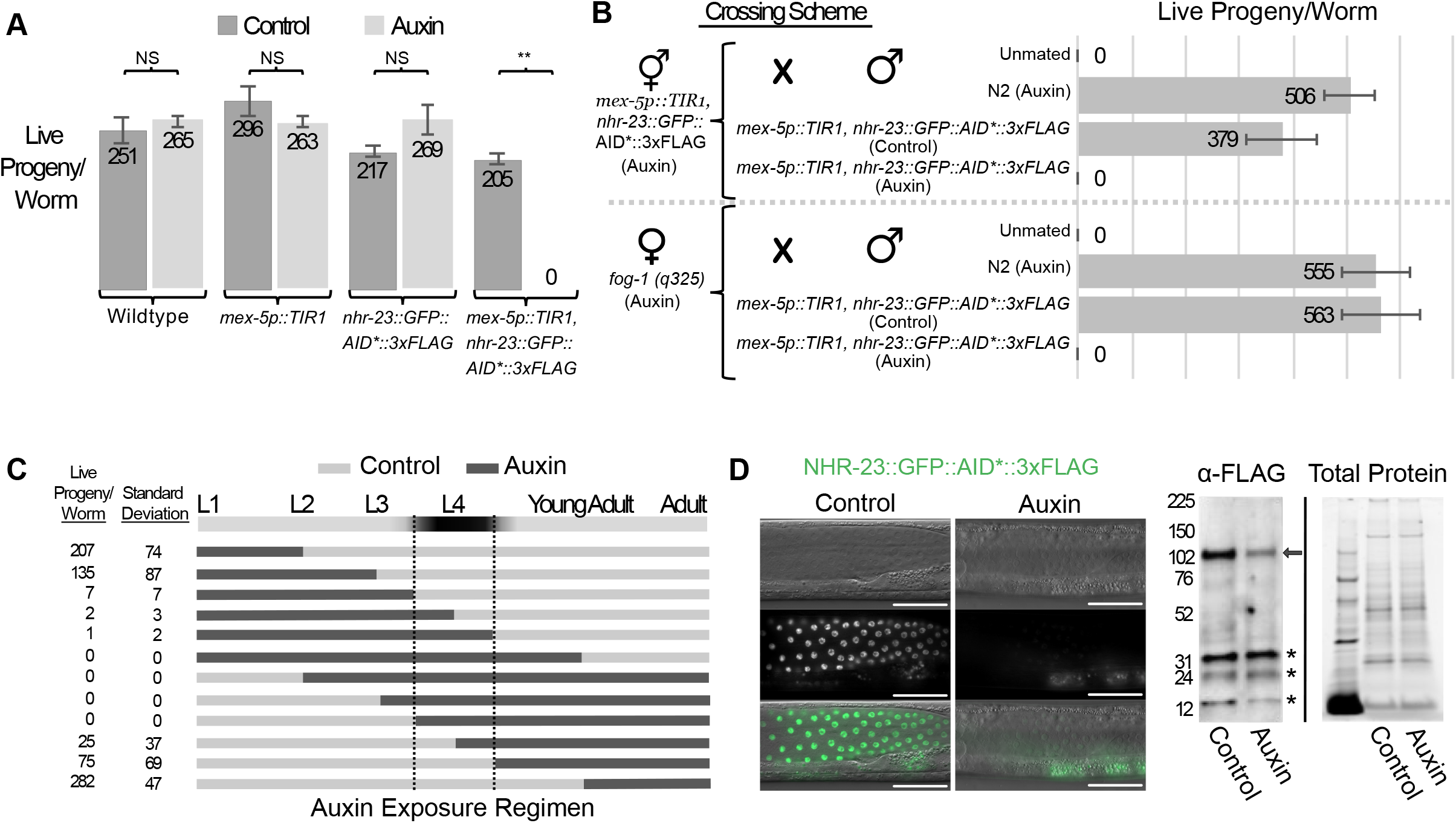
NHR-23-depletion in the germline during periods of spermatogenesis causes infertility in hermaphrodites and males. (A) Average number of live progeny produced by wildtype, *mex-5p:: TIR1, nhr-23::GFP::AID*::3xFLAG* or *mex-5p::TIR1 nhr-23::GFP::AID*::3xFLAG* hermaphrodites grown from L1 through adulthood on MYOB media with or without 4 mM auxin. A Student’s t-test was performed for each genotype comparing brood sizes on control media versus 4 mM auxin. A p-value greater than 0.01 was considered non-significant (NS). The **denotes p<0.00001. n=12 for each brood size. (B) Average number of live progeny produced by *mex-5p::TIR1 nhr-23::GFP::AID*::3xFLAG* or *fog-1(q325)* hermaphrodites grown from L1 on MYOB media with 4mM auxin. Hermaphrodites were either unmated or crossed to males of the indicated genotype; males were grown from L1 to adulthood on control or 4 mM auxin media. (n=12) (C) Average number of live progeny produced by *mex-5p:: TIR1 nhr-23::GFP::AID*::3xFLAG* hermaphrodites shifted on or off 4 mM auxin at different points in development. Dark horizontal bands represent growth on media with 4 mM auxin and light horizontal bands represent growth on control media lacking auxin. n=12 for each condition. (D) DIC and fluorescent images of *mex-5p::TIR1 nhr-23::GFP::AID*::3xFLAG* adult male germlines after animals were grown from L1 on MYOB media ±4 mM auxin. NHR-23-depletion results in minimally detectable GFP signal by fluorescence microscopy, but FLAG signal remains detectable in western blots. Scale bars: 40um. Anti-FLAG immunoblot analyses of lysates are from synchronized male *mex-5p::TIR1 nhr-23::GFP::AID*::3xFLAG* animals grown on control or 4 mM auxin media. Marker size (in kilodaltons) is provided. Arrowhead indicates NHR-23 fusion protein. Predicted size of the fusion protein is 86 kDa, based on the NHR-23 isoform expressed in adults in a Nanopore direct RNA sequencing dataset (Roach et al., 2020). Stain-free (Bio-Rad) analysis, which visualizes total protein on the membrane is provided as a loading control. The asterisks are non-specific bands previously observed with the anti-FLAG-HRP antibody (Ward, 2015).

We next carried out a set of experiments to determine when during development NHR-23 is necessary for fertility, akin to classic temperature-sensitive allele shifts to permissive/non-permissive temperatures. L1 synchronized *mex-5::TIR, nhr-23::GFP::AID*::3xFLAG* hermaphrodites were placed on control or 4 mM auxin media (Fig. 3C). Hermaphrodites were shifted from control media to 4 mM auxin, or vice versa, at the indicated timepoints. A short auxin exposure early in larval development (L1-L2) or continuous exposure after young adulthood had little or no effect on hermaphrodite self-fertility. The auxin-sensitive period was the L4 larval stage, with auxin exposure causing partially penetrant or complete self-infertility. Self-infertility was more pronounced in “shift-off” treatments, presumably due to the reported slow recovery of AID-tagged proteins following auxin removal (Zhang et al., 2015). Western blotting experiments revealed ~70% depletion of NHR-23 (Fig. 3D.)

While spermatogenesis is similar in *C. elegans* males and hermaphrodites, sex-specific differences do exist (L’Hernault et al., 1988; Minniti et al., 1996; Nance et al., 1999; Nance et al., 2000; Shakes and Ward, 1989). To test whether NHR-23 also functioned in male spermatogenesis, we repeated the crossing experiment from Fig. 3B using *mex-5::TIR1, nhr-23::GFP::AID*::3xFLAG* males. Males grown on control media sired progeny when crossed to either NHR-23-depleted hermaphrodites or feminized but otherwise wildtype *fog-1* “females”, but males grown on 4 mM auxin did not (Fig. 3B). Together, these data indicate NHR-23 is necessary for normal sperm production in both males and hermaphrodites.

### NHR-23 acts following the sperm/oocyte decision

The infertility phenotype could suggest that either NHR-23 was a spermatocyte/oocyte cell fate determinant or was responsible for executing the sperm fate program. To place NHR-23 in relation to known sex determination factors, we performed epistasis experiments (Fig. 4A). We crossed *nhr-23::GFP::AID*::3xFLAG* into a temperature sensitive (ts) gain-of-function *fem-3(q20)* allele genetic background (Barton et al., 1987). At 15°C, hermaphrodites produce both sperm and oocytes and NHR-23 expression was limited to the L4 stage (Fig. 4B). At 25°C, the masculinized germline produced only sperm, and NHR-23 expression persisted throughout adulthood. These results indicate NHR-23 functions downstream of FEM-3. Next, we tested *fog-3* (*feminization of germline*), which along with *fog-1* is believed to act as the terminal, germline-specific selector for sperm fate. As *fog-3* is close to *nhr-23* on chromosome I, we directly engineered a putative null allele of *fog-3* in an *nhr-23::GFP::BioTag::AID*::3xFLAG* strain (*wrd1* allele) by introducing a premature termination codon followed by a frameshift at the *fog-3* locus (*wrd6* allele). Because this experiment utilized a *nhr-23* strain distinct from the *wrd8* allele used elsewhere in this study, we verified that germline-specific NHR-23-depletion in a *sun-1p::TIR1; nhr-23(wrd1[GFP::BioTag::AID*::3xFLAG])* strain also caused sterility (Fig. S2). Importantly, this result demonstrates an equivalent phenotype with two distinct AID*-tagged *nhr-23* alleles and TIR1 transgenes being driven by two different germline-specific promoters (*mex-5p* vs *sun-1p*) (Fig. 3A; Fig. S2). In *fog-3(wrd6)* heterozygotes, NHR-23::GFP was expressed specifically in L4 hermaphrodites (Fig. 4C). In *fog-3(wrd6)* homozygotes, germline feminization is accompanied by a complete loss of NHR-23::GFP expression in L4 animals. These results show NHR-23 operates downstream of the known sex determination pathway, after spermatocyte cell fate has been decided.

**Fig. 4.**
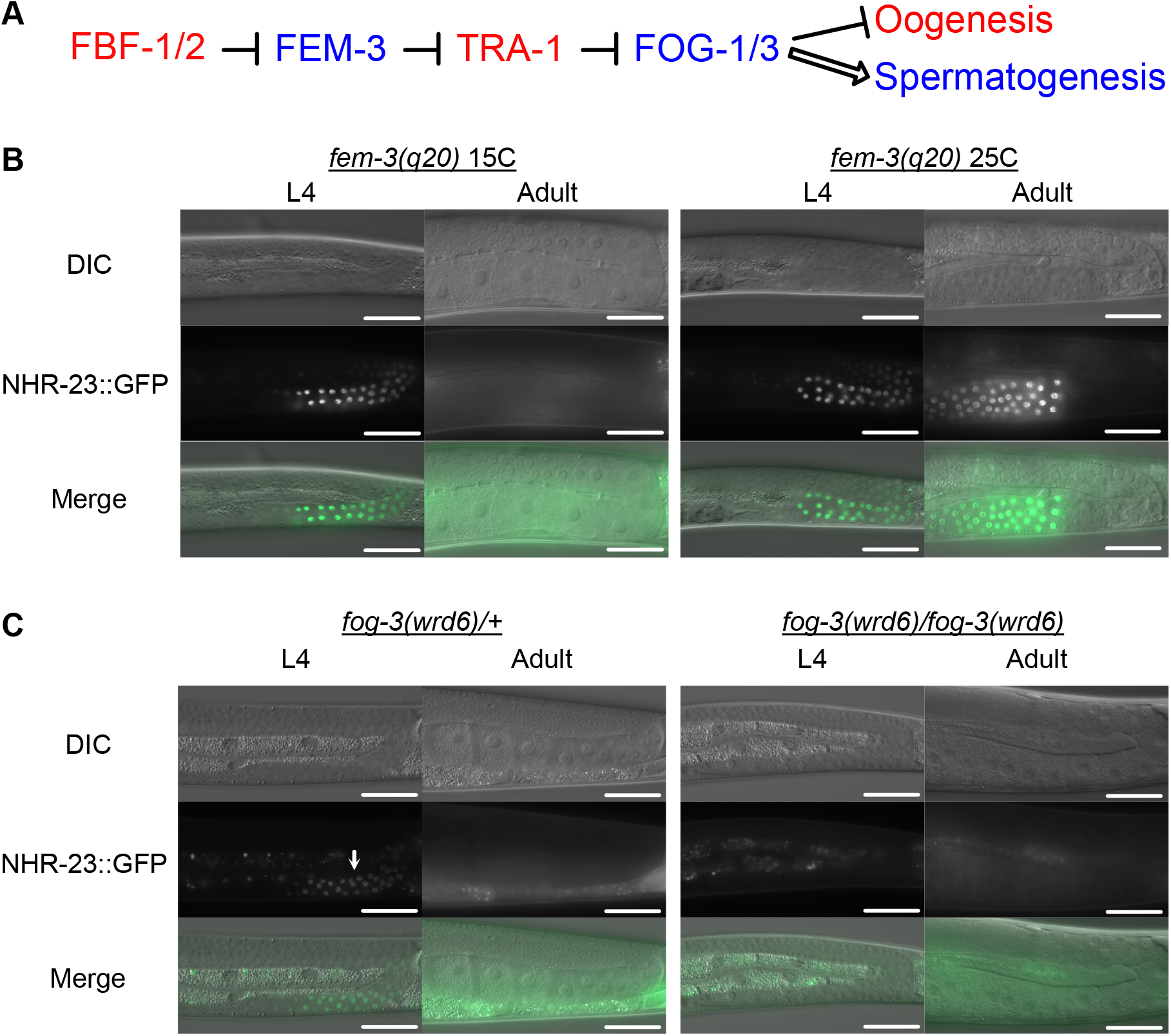
*nhr-23* is downstream of the germline sex determination pathway. (A) Simplified schematic of the *C. elegans* sex determination pathway and their effect on gamete fate. The wildtype function of the factors in blue text is to promote spermatogenesis, while the factors in red promote oogenesis. (B) DIC and GFP images of *mex-5p::TIR1 nhr-23::GFP::AID*::3xFLAG; fem-3(q20)* L4 and adult hermaphrodites grown from L1 onwards at permissive (15°C) or restrictive (25°C) temperatures. (C) DIC and GFP images of *mex-5p::TIR1 nhr-23::GFP::AID*::3xFLAG; fog-3(wrd6)* L4 and adult hermaphrodites. The white arrow indicates NHR-23::GFP::AID*::3xFLAG expression in late pachytene cells of a *fog-3(wrd6)* heterozygous hermaphrodite germline. Scale bars: 40 μm.

### Germline depletion of NHR-23 results in spermatocyte arrest

To better understand the nature of sperm-related fertility defects, we used DIC optics to examine gonads of control and NHR-23-depleted animals. The proximal gonad of control males contained a linear sequence of (1) spermatocytes completing meiotic prophase, (2) a discrete zone of meiotically dividing spermatocytes, budding figures, and residual bodies, and finally, (3) a zone of tightly packed spermatids with their small, highly refractive chromatin mass (Figs. 1B, 5A). NHR-23-depleted germlines produced spermatocytes that progressed normally through meiotic prophase. However, the rest of the gonad filled with what appeared to be primary spermatocytes, many of which contained small vacuoles. Haploid spermatids were notably absent. We observed similar defects when NHR-23 was depleted in masculinized germlines of *fbf-1(ok91) fbf-2(q704)* hermaphrodites (Fig. 5B). These results indicate NHR-23-depletion during male or hermaphrodite spermatogenesis results in a spermatocyte arrest phenotype.

**Fig. 5.**
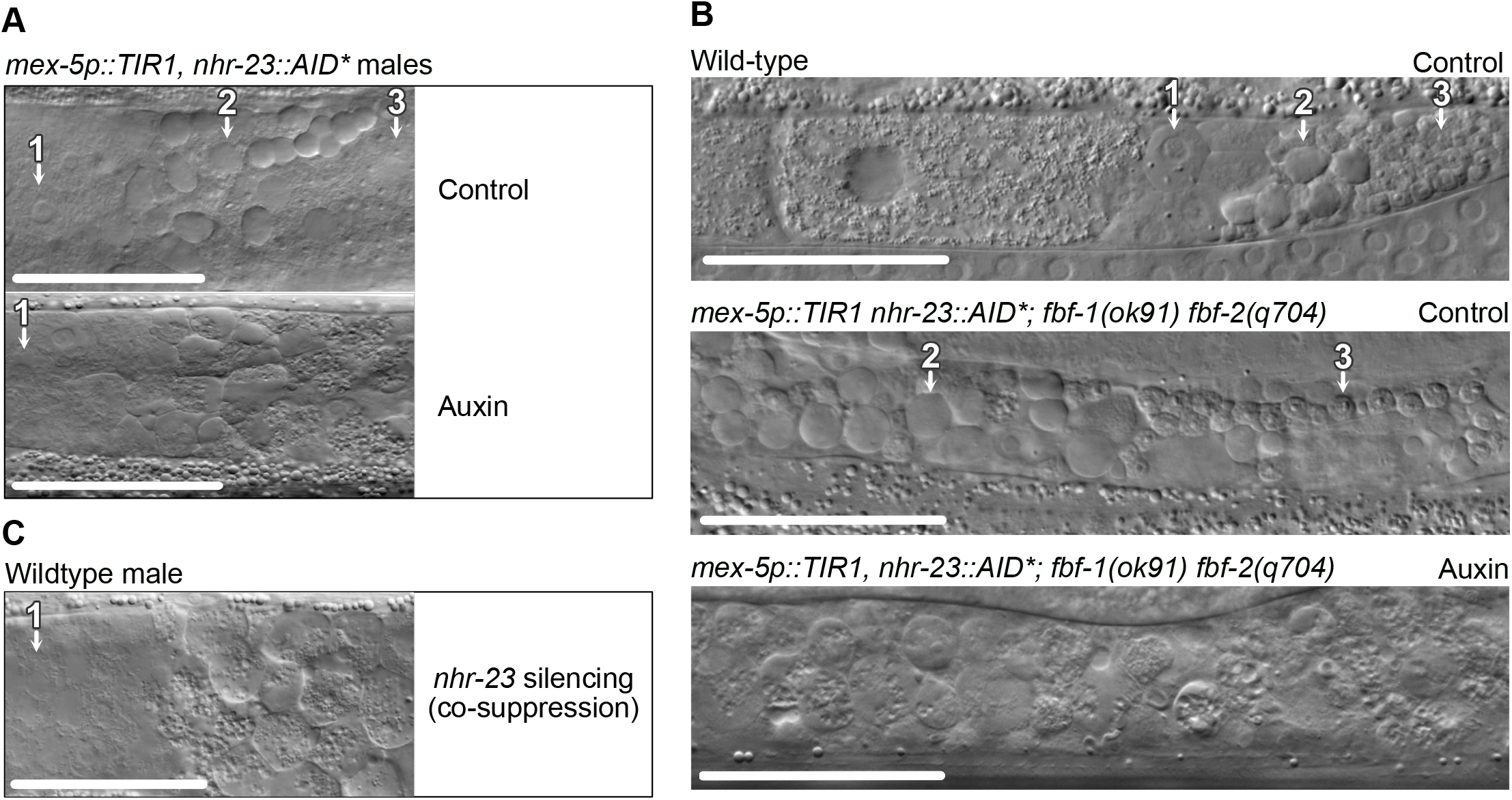
NHR-23 is necessary for sperm development. (A) DIC images of the proximal germlines of *mex-5p::TIR1 nhr-23::GFP::AID*::3xFLAG* young adult males grown from L1 onwards on control or 4 mM auxin media. (B) DIC images of wild type and masculinized *mex-5p::TIR1 nhr-23::GFP::AID*::3xFLAG fbf-1(ok91) fbf-2(q704)* hermaphrodites grown from L1 onwards on control or 4 mM auxin media. Primary spermatocytes (1) have a flat morphology with a large nucleus. Residual bodies (2) appear as highly refractile raised button-like structures. Spermatids (3) are readily discernible through their characteristic small refractive nuclei. (C) DIC image of the proximal germline in a wildtype young adult male F1 progeny from a P_0_ animal injected with a *mex-5p::nhr-23 cDNA* co-suppression construct, which promotes an RNAi-like depletion of targeted mRNA. Scale bars: 40 μm.

Because all experiments to this point relied on NHR-23-depletion via the AID system, we wanted to confirm this spermatocyte arrest phenotype through an independent method. We used transgene-mediated co-suppression, which has similarities to RNAi (Dernburg et al., 2000), to deplete *nhr-23* mRNA in the germline, and recovered viable but infertile males. These animals exhibited the same spermatogenesis defects as their auxin-mediated depletion counterparts, confirming the phenotypes are due to NHR-23-depletion (Fig. 5C).

### NHR-23-depleted primary spermatocytes arrest in a metaphase I-like state

We next wished to determine at which stages of spermatogenesis NHR-23 acts, so we assessed nuclear morphology in male germlines. Hoechst staining of DNA in control male germlines revealed the expected stages: i) diakinesis stage (Diak) spermatocytes with intact nuclear envelopes; ii) meiotically dividing spermatocytes; and iii) cells undergoing the post-meiotic partitioning (Par) event that yields a central residual body (RB) and four haploid spermatids (S) (Fig. 1, Fig. 6A). NHR-23-depleted male germlines included diakinesis spermatocytes and spermatocytes with chromosome patterns suggesting they were in prometaphase or metaphase I (Fig. 6B). Notably, these germlines lacked secondary spermatocytes, budding figures, or spermatids (Figs. 1D, 6B). In many cells, chromosomes were spread out relative to the initial metaphase clusters (Fig. 6B, early arrest), but there was no evidence of actual anaphase chromosome segregation, suggesting they had not moved beyond prometaphase or metaphase I. Only the oldest spermatocytes (farthest from the syncytial germline) contained the vacuoles observed in whole gonads (Fig. 5; late arrest). Nuclear envelope breakdown was independently confirmed by labelling fixed gonads with nuclear pore antibodies (Fig. 6J, unpublished data). These data suggested that spermatocyte development was aberrant, with an arrest in prometaphase or metaphase I.

**Figure 6.**
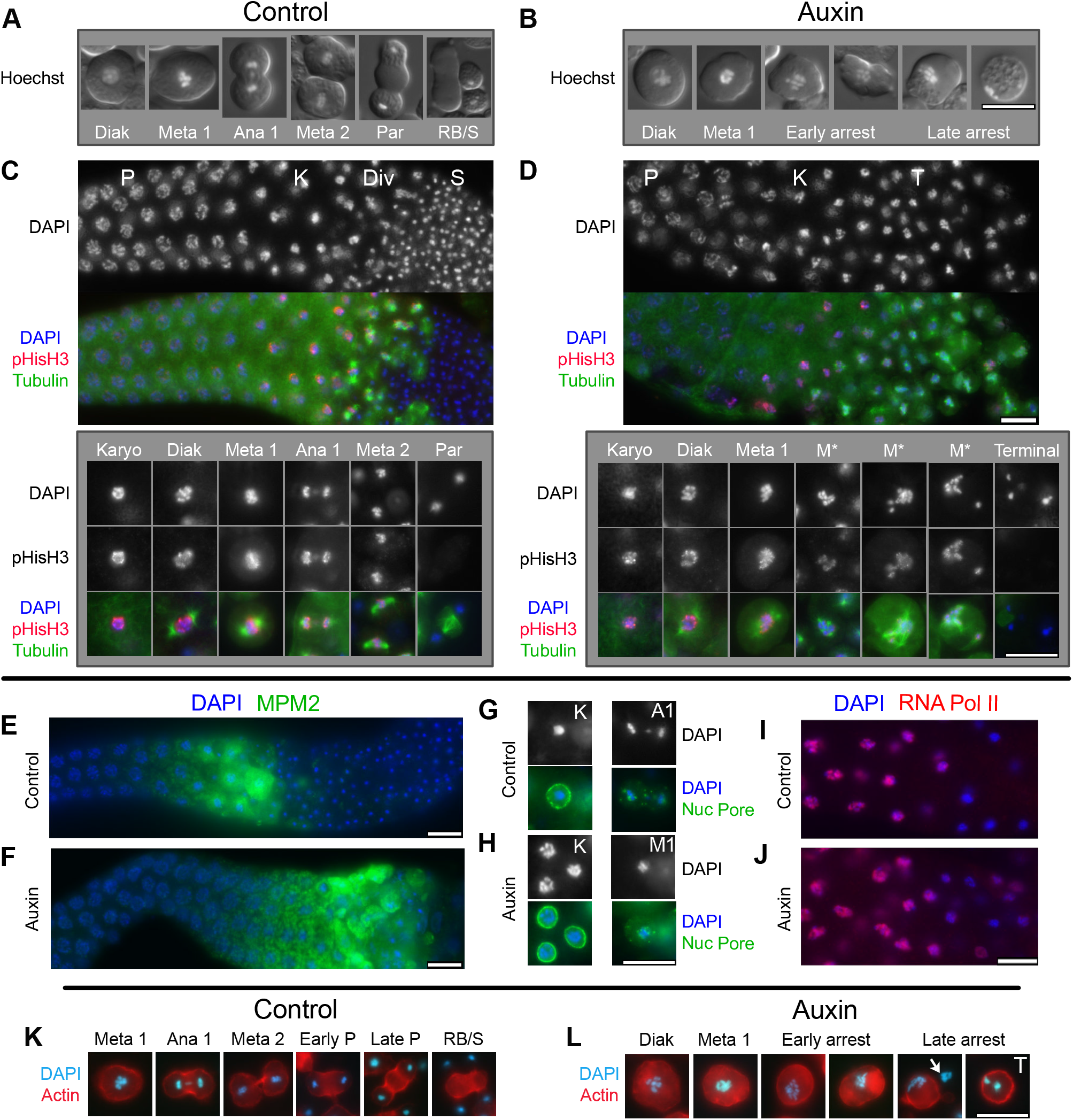
Metaphase I like arrest in NHR-23-depleted spermatocytes. (A,B) Unfixed spermatocytes ordered according to stage. Cells imaged with differential interference contrast (DIC), and DNA imaged with Hoechst stain. (C-J) Isolated and fixed male gonads and individual spermatocytes labeled with DAPI (blue) and indicated antibodies. (C,D) Gonad images show proximal gonad from late pachytene (P) to either haploid spermatids (S) or terminal arrest (T) co-labelled with antibodies against α-tubulin (green) and phosphorylated histone H3 (ser10) (red). Controls (*nhr-23*::degron without auxin or *him-5* with auxin) on left, and *nhr-23*::degron with auxin on right. Higher resolution images of individual spermatocytes below. (E-F). Isolated control (E) and NHR-23-depleted (F) proximal male gonads (pachytene and beyond) co-labelled with antibodies against MPM2 (green) which binds diverse mitotic and meiotic phosphorylated proteins. (G-H). Staged spermatocytes co-labelled with anti-nuclear pore protein (green) in control (G) and NHR-23-depleted (H) males. (I-J) Control (I) and NHR-23-depleted (J) gonads (pachytene through karyosome) co-labelled with anti-phospho-RNA polymerase II CTD repeat (red) shows turn-off of global transcription in karyosome spermatocytes. (K-L) Aldehyde fixed and staged spermatocytes with DNA pseudo-colored in cyan (DAPI) and actin microfilaments labeled with rhodamine-phalloidin (red). Arrow in late arrested image shows chromatin from lysed spermatocyte. Abbreviations: Pachytene (P), Karyosome (K), Meiotic Divisions (Div), Diakinesis (Diak), Metaphase I (Meta 1, MI), Aberrant Metaphase I (M*), Anaphase I (Ana 1, AI), Metaphase II (Meta 2), Post-meiotic Partitioning (Par), Residual Body (RB), Haploid Spermatids (S), Terminal-stage Spermatocyte (T). Scale bars: 10 μm.

To determine how well NHR-23-depleted spermatocytes were entering or exiting M-phase, isolated male gonads were first co-labelled with DAPI, anti-tubulin and anti-phosphohistone H3 Serine 10 (pHisH3(ser10)), a proxy for activity of the cell cycle regulator Aurora kinase (Hsu et al., 2000)(Fig. 6C,D). The most obvious NHR-23-depletion phenotype was an increase in the meiotic division zone (Div); large numbers of detached spermatocytes containing metaphase-like chromosomes with associated microtubule spindles (Fig. 6D). Recently detached spermatocytes had relatively normal diakinesis and metaphase I spindles, but metaphase chromosome alignment was impaired. Slightly older spermatocytes had more dispersed chromosomes (M*) and multipolar asters with four or more asters (Fig. 6D). In control germlines, anti-pHisH3(ser10) labelled chromosomes through both meiotic divisions and only became undetectable in post-meiotic budding figures. In NHR-23-depleted germlines, pHisH3 labelling persisted in arrested, metaphase-like cells. The very oldest spermatocytes exhibited a distinct terminal/late arrest phenotype; these spermatocytes lacked microtubule spindles and their chromosomes had aggregated into one or more tight masses and failed to label with anti-pHis3(ser10). To assess CDK-cyclin B activity, gonads were labelled for a phospho-epitope primarily found in proteins phosphorylated at inception of mitosis or meiosis (MPM2). In control spermatocytes (Fig. 6E), MPM2 labelling turned on during late meiotic prophase (karyosome stage), remained high through meiotic divisions, and then dropped abruptly during post-meiotic partitioning. In NHR-23-depleted spermatocytes (Fig. 6F), MPM2 labelling remained high in metaphase-arrested spermatocytes, only dropping in terminal arrest spermatocytes. Anti-nuclear pore antibodies confirmed that nuclear envelope breakdown occurred in a timely fashion in both control and NHR-23-depleted spermatocytes (Fig. 6G,H). Together, these results suggest that the cell cycle regulators of M-phase entry are unaffected by NHR-23-depletion. However depleted spermatocytes remain in a prolonged prometaphase state, develop multipolar spindles and never progress to anaphase. M-phase markers do eventually turn-off, but only in late, terminal stage spermatocytes.

We hypothesized that the observed meiotic defects might arise from earlier defects during the poorly understood karyosome stage (Fig. 1)(Shakes et al., 2009). In control gonads labelled with an antibody against active RNA polymerase II, entry into the karyosome stage was marked by abrupt cessation of global transcription (Fig 6I) (Shakes et al., 2009). In NHR-23-depleted gonads, turn-off of global transcription was both delayed and more gradual (Fig 6J). In a complementary fashion, the turn-on of pHisH3(ser10) labelling, which corresponds to chromosome condensation, was also delayed and more gradual (Fig. 6D). Closer examination revealed that, during the karyosome stage, the chromosomes in both control and depleted spermatocytes properly detached from the nuclear envelope, but the compaction of chromosomes into a single tight mass was more limited in NHR-23-depleted spermatocytes (Fig. 6G,H). Such aberrations in chromosome compaction during karyosome stage may contribute to subsequent defects in meiotic chromosome segregation as well as the prolonged M-phase arrest, if the defects trigger a cell cycle checkpoint.

### NHR-23-depleted spermatocytes have defects in both cytokinesis and post-meiotic partitioning

In previous studies, a metaphase I arrest of the cell cycle was not sufficient to block other aspects of sperm development. Notably, spermatocytes lacking a functional anaphase promoting complex arrest in metaphase I but still undergo post-meiotic partitioning to generate motile, albeit anucleate, sperm (Golden et al., 2000; Sadler and Shakes, 2000). Although NHR-23-depleted germlines do not make mature spermatids, we wondered if they nevertheless attempted aspects of normal, post-meiotic partitioning and residual body formation which is mediated in part by actin microfilaments (Hu et al., 2019; Winter et al., 2017). In control spermatocytes, actin localization to the cortex was uniform in metaphase and enriched in cortical rings during anaphase (Fig. 6K). In partitioning stage spermatocytes, actin localized in a central band and was simultaneously lost from the cortex of budding spermatids (Fig. 6K). In NHR-23-depleted spermatocytes, actin patterns remained largely cortical in both metaphase arrested and late terminally arrested spermatocytes (Fig. 6L). Consistent with our DIC analysis, NHR-23-depleted spermatocytes showed no evidence of cytokinesis or residual body formation. These results are consistent with NHR-23-depletion causing defects in addition to meiotic chromosome segregation.

### NHR-23 is required for biogenesis of sperm-specific fibrous body-membranous organelle (FB-MO) complexes

Among *C. elegans* mutants with spermatogenesis defects, several exhibit a spermatocyte-arrest phenotype. Four of these (*spe-4, spe-5, spe-6*, and *wee-1.3*) exhibit a vacuolated phenotype by DIC very similar to NHR-23-depleted spermatocytes (Fig. 7A). Other spermatocyte arrest mutants (spe-39, *spe-44*) do not exhibit the same vacuolated phenotype (Fig. 7A)(Kulkarni et al., 2012; Zhu and L’Hernault, 2003). In previous studies of vacuolated mutants, transmission electron microscopy found these vacuoles are swollen, distended membranous organelles (MOs) (Lamitina and L’Hernault, 2002; L’Hernault and Arduengo, 1992; Machaca and L’Hernault, 1997; Varkey et al., 1993). To determine if vacuoles in NHR-23-depleted spermatocytes were MOs, we crossed a PEEL-1::GFP transgene into our *mex-5p::TIR1, nhr-23::GFP::AID*::3xFLAG* strain. PEEL-1 is the toxin component of a toxin-antidote selfish genetic element (Seidel et al., 2008) that also serves as a useful MO marker (Seidel et al., 2011). In control primary spermatocytes and budding figures, PEEL-1::GFP marked internal membranes (Figs 1 and 7B). In mature spermatids, PEEL-1::GFP exhibited a bright punctate pattern corresponding to plasma membrane-docked MOs. In NHR-23-depleted spermatocytes, PEEL-1::GFP colocalized with vacuoles in the corresponding DIC images (Fig. 7B), indicating that MOs form but then vacuolate in terminal arrest spermatocytes.

**Figure 7.**
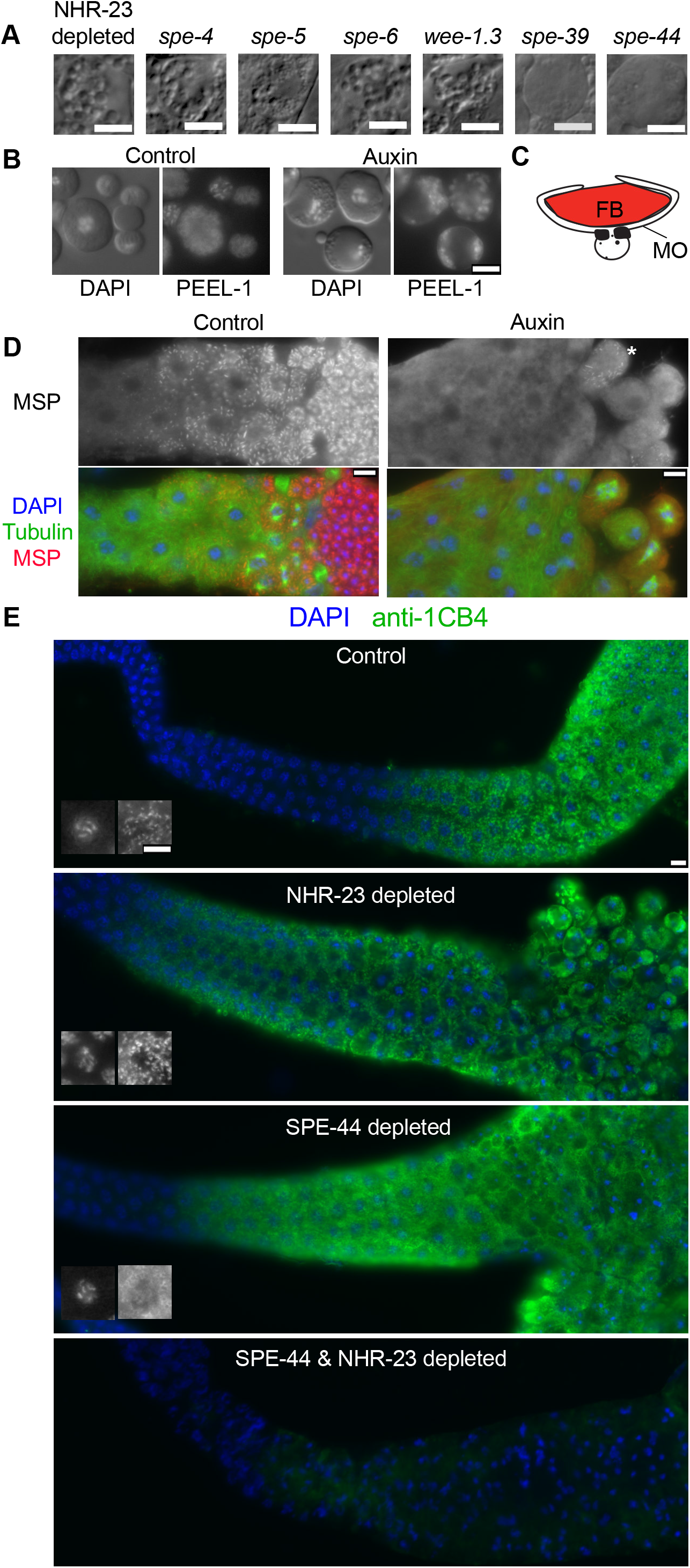
FB-MO defects in NHR-23 spermatocytes and synergy with SPE-44. (A) DIC images of arrested spermatocytes from spermatogenesis-defective mutants. (B) Control and NHR-23-depleted spermatocytes visualized by DIC/Hoechst (left) and the MO marker PEEL::GFP (right). (C) Schematic of an FB-MO complex showing an MSP-enriched fibrous body (FB) enveloped within arm-like extensions of the Golgi-derived membranous organelle (MO). (D) Proximal gonads from control and NHR-23-depleted males co-labelled with DAPI and antibodies against tubulin and MSP. Asterisk indicates a single spermatocyte in the NHR-23-depleted gonad with MSP polymers. (E) Gonads from animals with indicated protein depletions co-labelled with DAPI and the anti-MO antibody 1CB4. Large DAPI and 1CB4 inserts show MO patterns in pachytene stage spermatocytes and highlight defects in SPE-44 spermatocytes. Scale bars: 5 μm.

In developing spermatocytes, MOs serve as sites where Major Sperm Protein (MSP) assembles into fibrous bodies (FBs) (Fig. 7C). MSP is a nematode-specific protein which drives the actin-like cell motility of spermatozoa while also serving as a signaling molecule for oocyte maturation (Miller et al., 2001). Not surprisingly, MO defects are often associated with either abnormalities in FB assembly as in *spe-5* and *spe-39* mutants (Machaca and L’Hernault, 1997; Zhu and L’Hernault, 2003) or a complete failure in FB formation as in *spe-6* and *spe-44* (Kulkarni et al., 2012; Varkey et al., 1993). In wildtype animals, MSP is first expressed in mid-pachytene spermatocytes and, by the end of the karyosome stage, individual spermatocytes fill with discrete, uniformly sized FBs (Fig. 7D)(Chu and Shakes, 2013). During post-meiotic partitioning, FB-MO complexes facilitate MSP delivery to spermatids. In NHR-23-depleted spermatocytes, MSP was expressed, but in most spermatocytes, it remained dispersed throughout the cytoplasm (Fig. 7D). NHR-23-depleted germlines sometimes included individual spermatocytes with MSP assemblages (Fig. 7D, asterisk) that were more variable in shape and size than normal FBs.

To investigate potential defects in FB-MO biogenesis, gonads were immunolabelled with 1CB4, an antibody labelling glycosylated MO proteins (Okamoto and Thomson, 1985). In control gonads, 1CB4 labelled discrete cytoplasmic organelles starting in mid-pachytene spermatocytes (Fig. 7E). This pattern persisted until the spermatid stage when individual MOs docked with the plasma membrane (Fig. 1D, 7E). In NHR-23-depleted germlines, the 1CB4 pattern in pachytene spermatocytes was similar to controls. However, in arrested 1° spermatocytes, MOs aggregated into clumps adjacent to the plasma membrane, suggesting a defect in either MO docking or later stages of MO morphogenesis.

Of spermatocyte arrest mutants shown in Fig. 7A, only the *spe-44* gene encodes an early transcriptional regulator (Kulkarni et al., 2012). Like NHR-23-depleted spermatocytes, *spe-44* spermatocytes exhibit defects in MSP assembly and meiotic progression (Kulkarni et al, 2012). However *spe-44* spermatocytes arrest in the second rather than the first meiotic division, and arrested spermatocytes do not fill with distended MOs (Fig. 7A). To determine whether NHR-23 and SPE-44 play similar or distinct roles in spermatogenesis, we compared 1CB4 patterns as a proxy for their role in FB-MO biogenesis. In SPE-44-depleted pachytene spermatocytes, the 1CB4 pattern within developing spermatocytes did not form discrete structures (Fig. 7E). This 1CB4 pattern matches that published for *spe-44(ok1400)* mutants (Kulkarni et al., 2012) and is distinct from NHR-23-depletion defects. As NHR-23 and SPE-44 individual depletions produce similar, but distinct MO defects, we simultaneously depleted both and found the double depletion had a strong synthetic effect and almost completely abolished 1CB4 staining (Fig. 7E).

### NHR-23 and SPE-44 independently regulate spermatogenesis

The synthetic FB-MO phenotype observed following NHR-23+SPE-44 double depletion raised the possibility that these regulators could function in distinct pathways. In *spe-44(ok1400)* mutants, levels of *nhr-23* mRNA do not differ significantly from wildtype (Kulkarni et al., 2012). To test whether SPE-44 might function downstream of NHR-23, we generated a strain carrying an *mScarlet::3xMyc* cassette inserted into the 3’ end of *spe-44* by CRISPR/Cas9-mediated genome editing. There is only a single *spe-44* isoform and our strain results in a C-terminal translational fusion. We crossed this *spe-44::mScarlet::3xMyc* allele into a strain carrying *mex-5::TIR1, nhr-23::GFP::AID*::3xFLAG*. On control media, spermatocytes in this strain exhibited an almost complete overlap in expression for SPE-44 and NHR-23 (Fig. 8A). When grown on auxin, depletion of NHR-23 did not affect the level or expression pattern of SPE-44::mScarlet::3xMyc. These data indicate NHR-23 regulates neither SPE-44 expression nor its localization.

**Fig. 8.**
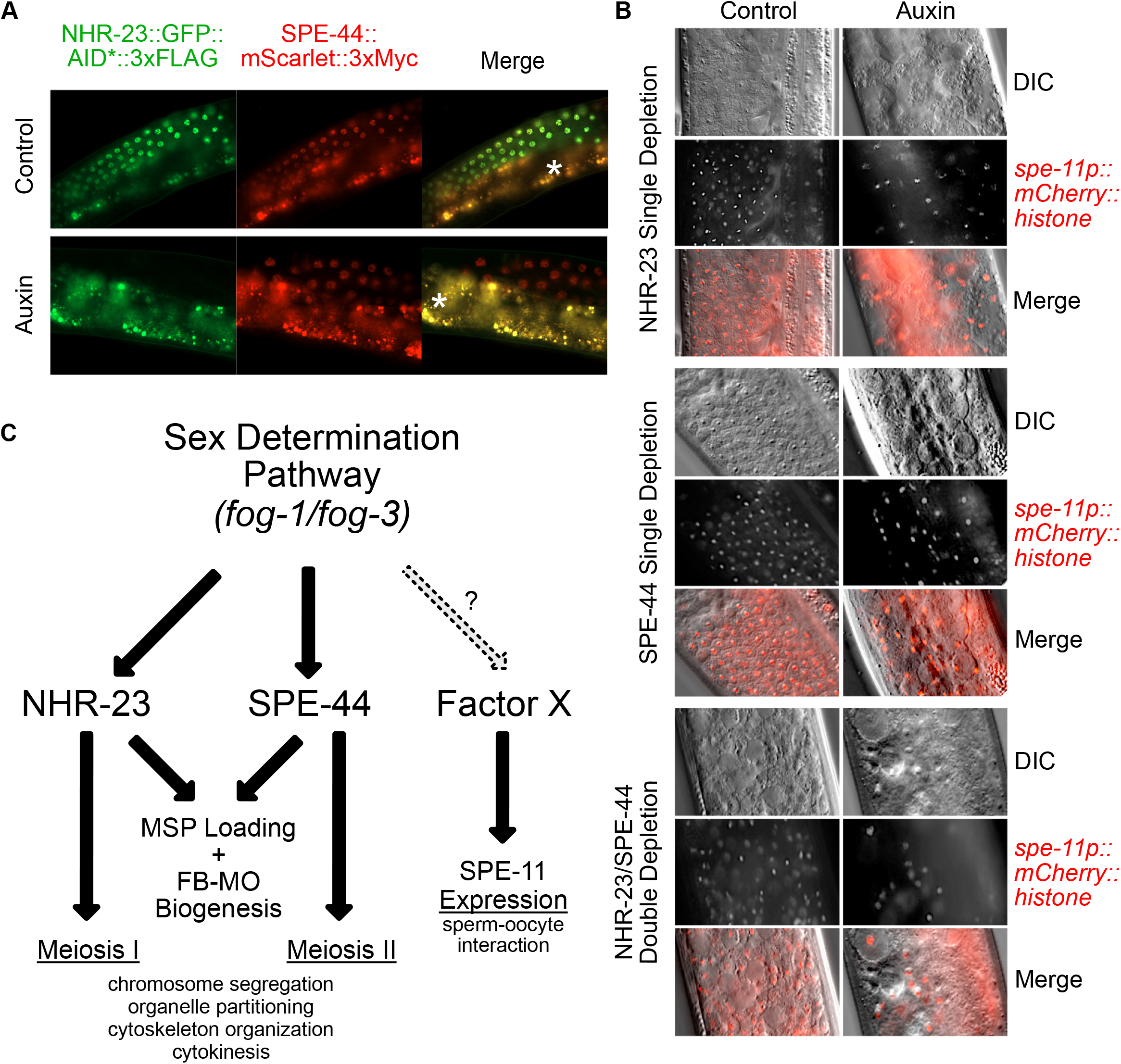
Control of transcription in spermatogenesis is facilitated by multiple factors with divergent targets. (A) Fluorescent images of NHR-23::GFP::AID*::3xFLAG and a mScarlet::3xMyc knock-in to the endogenous *spe-44* gene to create a C-terminal translational fusion. Worms containing both translational fusions as well as the *mex-5p::TIR1* driver were grown from L1 onwards on control or 4 mM auxin media. The asterisks indicate gut autofluorescence. (B) Representative DIC and fluorescent images of *spe-11p::mCherry::H2B* in adult males expressing *mex-5p::TIR1* in the germline. Worms individually expressed NHR-23::AID*::3xFLAG or SPE-44::AID*::3xFLAG or both and were grown from L1 until the young adult stage on control or 4 mM auxin media. (C) Model depicting the coordinated control of gene expression prior to and during the stages of spermatogenesis. NHR-23 and SPE-44 control distinct sets of genes to promote the events of meiosis I and II, respectively. They redundantly promote MSP loading and FB-MO biogenesis. A third pathway controlled by yet unidentified factors promotes the expression of genes involved in the sperm-oocyte interaction, such as *spe-11*.

These data raised the possibility that *C. elegans* spermatogenesis coordinated by multiple transcriptional regulatory pathways. If so, then spermatogenesis-specific genes like *spe-11* that are known not to be regulated by SPE-44 may instead be NHR-23 target genes. We acquired a strain in which the *spe-11* promoter drives expression of an *mCherry*::Histone *H2B::unc-54 3’UTR* reporter (Frøkjaer-Jensen et al., 2008). We crossed AID* alleles of *nhr-23* (Zhang et al., 2015) and *spe-44* (Kasimatis et al., 2018) lacking a fluorescent protein fusion, as well as a *mex-5p::TIR1* allele, into this *spe-11* reporter strain. In animals grown on control media, the *spe-11* promoter drove mCherry-histone expression starting in pachytene spermatocytes with the histone fusion protein persisting in spermatids (Fig. 8B(Frøkjaer-Jensen et al., 2008). Following individual depletion of either NHR-23 or SPE-44, expressed mCherry fusion protein persisted in meiotically arrested nuclei despite the spermatocyte defects and their meiotic arrest in meiosis I or II, respectively. To test whether NHR-23 and SPE-44 redundantly regulated *spe-11*, we created a *spe-11* reporter strain that would allow simultaneous germline-specific depletion of NHR-23 and SPE-44. Co-depletion of NHR-23 and SPE-44 resulted in germline defects so severe that it was difficult to identify individual cells within the proximal gonad. Yet, reporter expression was still detectable in nuclei (Fig. 8B). Together these results suggest not only that NHR-23 and SPE-44 function in independent pathways, but also that *spe-11* is regulated either by factors upstream of NHR-23 and SPE-44 or by an additional, yet to be identified transcriptional regulatory pathway.

## Discussion

*C. elegans* spermatogenesis offers a powerful model to define how gene-regulatory networks implement and coordinate the cell cycle and developmental programs. In this study, we identify NHR-23 as a critical regulator of spermatogenesis in both hermaphrodite and male animals.

NHR-23 is expressed in 1° spermatocytes and acts in the L4 stage in hermaphrodites, equivalent to the temperature-sensitive (ts) period for ts alleles of diverse spermatogenesis regulators (Hirsh and Vanderslice, 1976; Varkey et al., 1995; Ward and Miwa, 1978). Upon inactivation of NHR-23 through either co-suppression-mediated mRNA silencing (Fig. 5) or auxin-induced protein degradation (Fig. 3–5), affected hermaphrodites failed to produce mature spermatids. NHR-23 functions downstream of the sex determination pathway (Fig. 4) as cells recognizable as 1° spermatocytes developed (Fig. 5–7) and progressed through meiotic prophase (Fig. 6). Numerous cell cycle markers (Aurora kinase and CDK-cyclinB activity proxies, nuclear envelope breakdown) indicated that affected spermatocytes entered meiosis but then failed to divide (Fig. 6). Instead, the fully condensed chromosomes clustered within a disorganized and ultimately multipolar microtubule array (Fig. 6). Although microtubules appeared to associate with the chromatin, there was no evidence of even attempted anaphase segregation (Fig. 6). Similarly, we never observed cytokinesis or spermatid budding divisions. The oldest cells contained terminal chromatin masses, which may represent either a variation of normal sperm chromatin remodeling or apoptotic compaction (Fig. 6). In addition to these meiotic roles, NHR-23 is necessary for proper FB-MO biogenesis. Golgi-derived MOs developed in meiotic prophase spermatocytes, yet MSP failed to assemble into associated FBs and the morphology of MOs in arrested spermatocytes was clearly abnormal (Fig. 7).

### Multiple gene regulatory networks may control *C. elegans* spermatogenesis

The meiotic phenotypes following NHR-23-depletion are similar to but distinct from those caused by inactivation of *spe-44*, a critical transcriptional regulator of spermatogenesis (Fig. 8C). In both NHR-23-depletion and *spe-44* mutants, meiotic prophase is largely unaffected except that chromatin compaction is delayed during the karyosome stage (Kulkarni et al., 2012). *spe-44* spermatocytes also fail to assemble MSP into FBs (Kulkarni et al., 2012). Both NHR-23 and SPE-44 proteins are first expressed in developing spermatocytes and have similar mRNA expression profiles in a spatiotemporal analysis of germline expression (Tzur et al., 2018). NHR-23 and SPE-44 both function downstream of the canonical sex determination pathway (Fig. 4) (Kulkarni et al., 2012). While both are essential to complete the spermatocyte meiotic divisions, NHR-23 is necessary for the first meiotic division (Fig. 6), whereas SPE-44 is necessary for the second meiotic division (Fig. 8C)(Kulkarni et al., 2012). Their microtubule spindle patterns also differ; in NHR-23-depleted spermatocytes the microtubule asters remain close to the chromosomes and cell center (Fig. 6D), whereas those in spe-44 mutants move to the cell cortex (Kulkarni et al., 2012) SPE-44 regulates ELT-1 (del Castillo-Olivares et al., 2009; Kulkarni et al., 2012), which promotes MSP gene expression in conjunction with a histone methyltransferase, SET-17 (Engert et al., 2018). However, SPE-44 does not regulate *nhr-23* (Kulkarni et al., 2012), nor does NHR-23-depletion affect SPE-44 expression (Fig. 8A) consistent with these two transcription factors regulating distinct pathways. Co-depletion of NHR-23 and SPE-44 led to additive phenotypes such as loss of the MO marker 1Cb4 (Fig. 7E) and more severe germline morphology defects (Fig. 8B). NHR-23-depleted spermatocytes developed distended MOs similar to those of several sperm function mutants (e.g. *spe-4(lf), spe-6(lf)*, *wee-1.3(gf)*)(Lamitina and L’Hernault, 2002; L’Hernault et al., 1988; Muhlrad and Ward, 2002; Varkey et al., 1993). *spe-44* does not regulate these genes (Kulkarni et al., 2012) and does not exhibit this mutant phenotype (Kulkarni et al., 2012) (Fig. 7A). These results suggest that NHR-23 and SPE-44 function in separate pathways to regulate distinct cellular processes.

Additional undiscovered gene regulatory pathways are likely to help execute the *C. elegans* spermatogenesis program. *spe-11p::mCherry::H2B* reporter activity was unaffected by NHR-23, SPE-44, or NHR-23+SPE-44 depletions. These data could suggest the presence of three or more distinct pathways controlling sperm morphogenesis in *C. elegans*, or alternatively *spe-11* is directly regulated by a factor upstream of both NHR-23 and SPE-44 (Fig. 8C). Notably, SPE-44 only accounts for one-quarter of sperm-expressed genes (Kulkarni et al., 2012). As we further characterize NHR-23 and SPE-44, it will be informative to compare their function to spermatogenesis regulators in other systems. In *Drosophila*, BAM promotes differentiation of spermatogonia into spermatocytes (McKearin and Spradling, 1990), while the sperm-specific transcriptional program is promoted by two complexes: tMAC and tTAF (Beall et al., 2007; Laktionov et al., 2018; Metcalf and Wassarman, 2007). In mice, retinoic acid receptor signaling promotes both spermatogonia differentiation and meiotic entry (Chung et al., 2004; Gely-Pernot et al., 2012; Gely-Pernot et al., 2015), and MYBL1 coordinates the expression of four downstream transcription factors (CREM-τ, TRF2, RFX2 and Sox30) (Bolcun-Filas et al., 2011; Horvath et al., 2009; Li et al., 2013; Zhang et al., 2018). Zhang *et al*. (2018) suggests that tree-like regulatory cascades would be more efficient than a network enriched in nodes. Spermatogenesis is a rapidly evolving process. Defining the regulatory architecture across a range of model organisms will shed insight into conserved features controlling sperm morphogenesis as well as distinctive features which have evolved separately in mammals, insects, nematodes, and other organisms (White-Cooper and Bausek, 2010).

### Circadian rhythm regulators in spermatogenic tissues

Previous to this study, the best characterized role for *nhr-23* in *C. elegans* was as a key regulator of molting (Frand et al., 2005; Kostrouchova et al., 1998; Kostrouchova et al., 2001; Kouns et al., 2011; Patel and Frand, 2018). Like many molting factors, *nhr-23* mRNA levels oscillate over the course of each larval stage (Gissendanner et al., 2004; Hendriks et al., 2014; Kostrouchova et al., 2001). Yet in sperm-producing germlines, NHR-23 is expressed constitutively, suggesting a radically different mode of regulation in soma vs. germ tissues. Interestingly, many core mammalian circadian rhythm regulators also oscillate in the soma but exhibit constitutive, non-oscillatory expression in the testes (Alvarez et al., 2008; Kang et al., 2010; Kennaway et al., 2012; Morse et al., 2003). Although the non-circadian roles of these factors in spermatogenesis have not been deeply explored, null male mice of the clock gene BMAL1 have fertility defects and are difficult to mate (Alvarez et al., 2008), and depletion of the NHR-23 homolog, RORα/NR1F1, in rat Sertoli cells reduces sperm count (Mandal et al., 2018). It remains unclear how these timers have been co-opted to promote spermatogenesis. Our work may provide an entry point into understanding both the function of these circadian rhythm factors during spermatogenesis and how their expression is altered from oscillating to constitutive expression in the testes.

### Future perspectives

Most known spermatogenesis regulators have been identified in forward genetic screens and candidate-based testing of male germline-enriched factors. These approaches have been incredibly powerful but would miss factors such as NHR-23 which also have earlier essential roles in embryogenesis and molting. Similarly, as NHR-23 is maternally loaded, it was not identified by focusing on male enriched transcriptional regulators (Ebbing et al., 2018; Kulkarni et al., 2012; Reinke et al., 2004; Tzur et al., 2018). High-throughput GFP::AID* tagging of NHR-23-regulated genes could provide an exciting avenue for identifying novel regulators of spermatogenesis. New datasets that detail mRNA expression with respect to position in the germline can also be analyzed to identify transcriptional regulators expressed before, coincident with, and after the NHR-23/SPE-44 zone of expression. Going forward, determining factors that either regulate or are regulated by NHR-23 and SPE-44 will deepen our understanding of how the NHR-23 node controls spermatogenesis and how gene regulatory networks control the complex cellular events underpinning sperm morphogenesis.

## Materials and Methods

### Strains and Culture

*C. elegans* were cultured as originally (Brenner, 1974), except worms were grown on MYOB media instead of NGM. Strains used in this study:

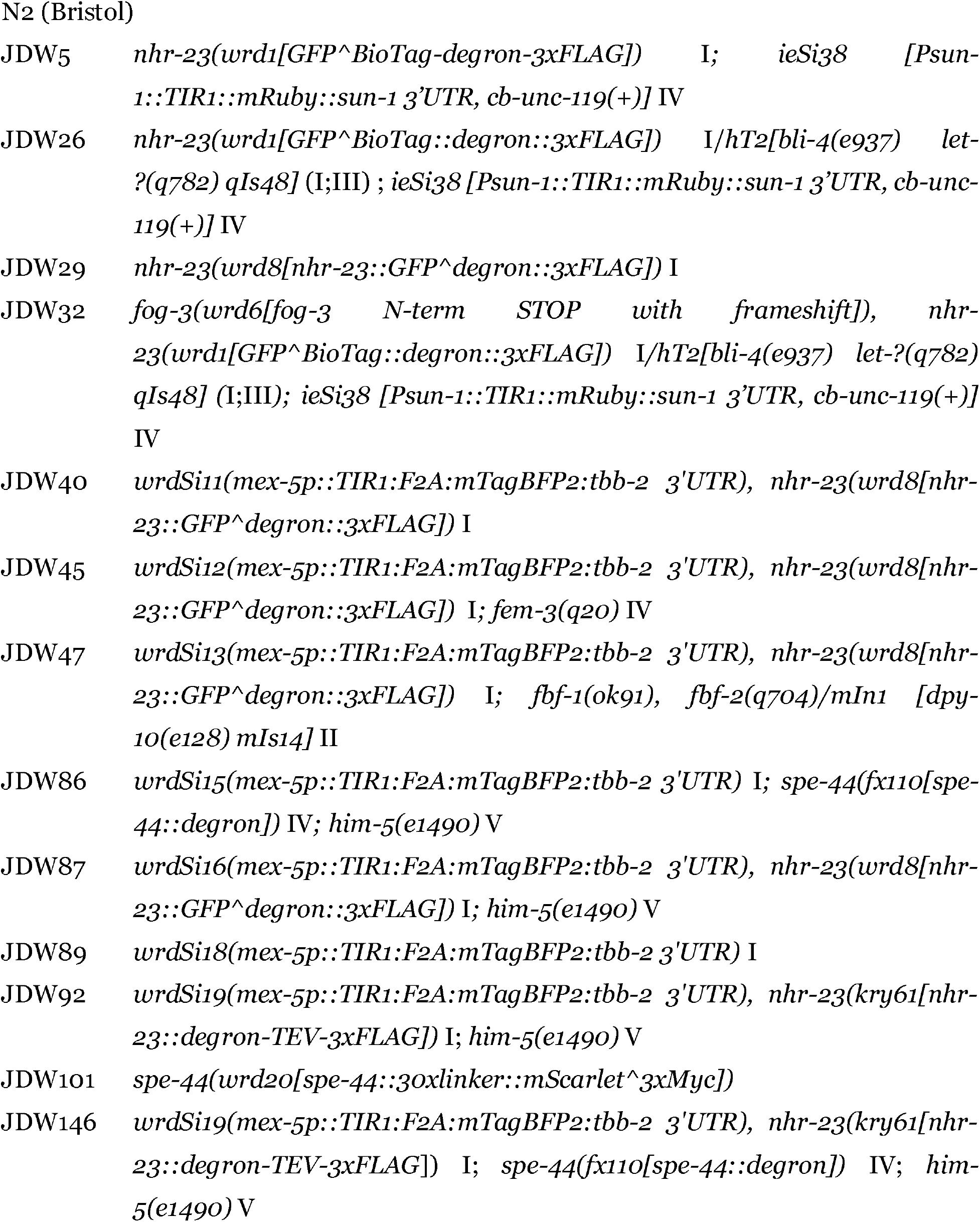
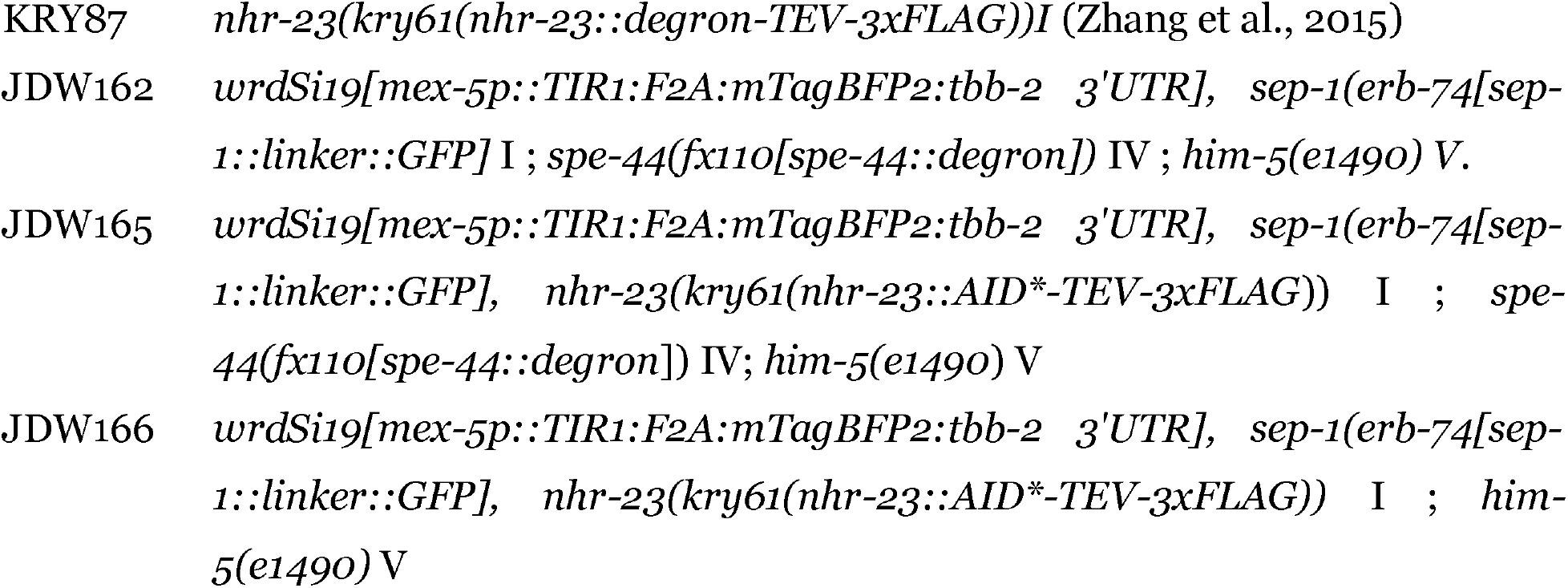

The following strains were provided by the CGC, which is funded by NIH Office of Research Infrastructure Programs (P40 OD010440):

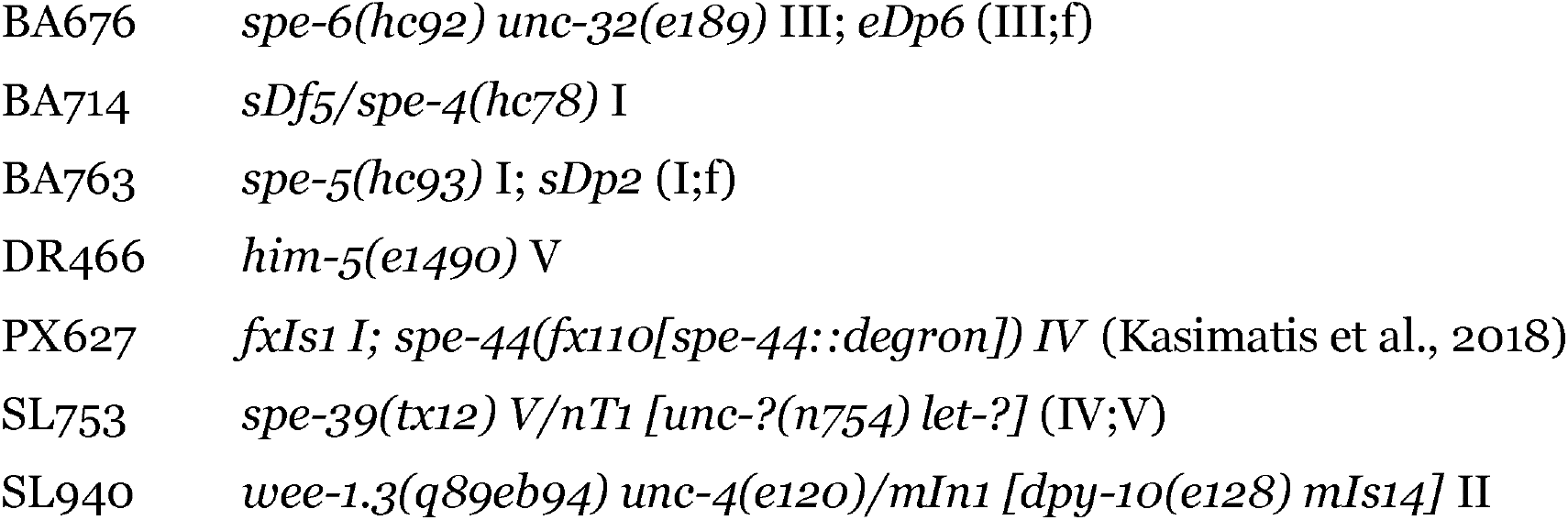

The following was a gift from Erik Jorgensen and is currently unpublished:

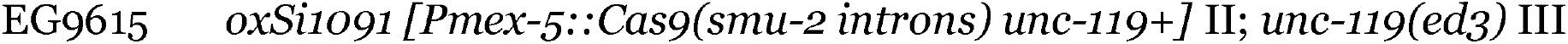

### Immunocytochemistry

Intact gonads were obtained by dissection of individual males in 5-10 microliters of egg buffer (Edgar, 1995) on ColorFrost Plus slides (Fisher Scientific #12-550) coated with poly-L-lysine (Sigma Aldrich #P8290). Samples were freeze-cracked in liquid nitrogen. Sperm spreads to analyze detached spermatocytes and spermatids were obtained by applying slight pressure to the coverslip before freeze-cracking. Most samples were fixed overnight in −20°C methanol. Specimen preparation and antibody labeling followed established protocols (Shakes et al., 2009). Primary antibodies included: 1:100 Alexa Fluor 488-conjugated anti-α-tubulin (mouse monoclonal DM1A, Millipore-Sigma #16-323), 1:200 rabbit anti-phosphorylated histone H3 (S10) (Millipore #06-570, Lot 2649123), 1:100 MPM-2 mouse monoclonal (EMD Millipore #05-368), 1:200 anti-nuclear pore antibody Mab414 (Abcam ab24609), 1:250 anti-RNA polymerase II CTD repeat Ser2 (Abcam #ab5095), 1:200 rabbit anti-SPE-44 (Kulkarni et al., 2012), 1:15,000 G3197 rabbit anti-MSP polyclonal (Kosinski, 2005; gift from David Greenstein, University of Minnesota), and 1:40 1CB4 monoclonal (Okamoto and Thomson, 1985; gift from Steve L’Hernault, Emory University). All samples were incubated with primary antibodies for 2 hours at room temperature except MPM-2 samples which were incubated overnight at 4°C. Affinity-purified secondary antibodies included 1:300 Alexa Fluor Plus 555 goat anti-rabbit IgG (Invitrogen #A32732) and 1:100 Alexa Fluor 488 Goat anti-mouse IgG (H+L) (Jackson ImmunoResearch #115-545-146).

Final slides were mounted with DAPI containing Fluoro Gel II mounting medium (Electron Microscopy Sciences #50-246-93). Images were acquired under differential interference contrast or epifluorescence using an Olympus BX60 microscope equipped with a QImaging EXi Aqua CCD camera. Photos were taken, merged, and exported for analysis using the program iVision. In some cases, the levels adjust function in Adobe Photoshop was used to spread the data containing regions of the image across the full range of tonalities.

For actin staining, slides with dissected gonads were fixed for 10 min in 4% fresh paraformaldehyde (FisherScientific) in 1x PBS. Samples were quenched in 1 M glycine (FisherScientific) in PBS (FisherScientific #BP3994) for at least 5 min, permeabilized in 0.1% Triton-X-100 (FisherScientific #BP151) in PBS for 5 min, then dip washed in 1xPBS. Slides were incubated with rhodamine-conjugated phalloidin (Invitrogen #R415) diluted 1:100 in 1xPBS for 15 min in the dark before washing three times in PBS for 5 min each. Slides were mounted as above.

For DIC/Hoechst preparations, males were dissected in buffer with 100 mg/ml Hoechst 33342 (Sigma Aldrich #94403) on non-plus slides and immediately imaged.

### CRISPR/Cas9 Genome Editing

A thorough description of the plasmids and oligonucleotides described in this section is provided in Table S1 and S2, respectively.

*nhr-23(wrd1[GFP^BioTag::degron::3xFLAG])* was generated using a pJW1598 repair template, based on the previously described set of self-excising cassette (SEC) vectors for genome editing (Dickinson et al., 2015). Briefly, *nhr-23* 5’ and 3’ homology arms were PCR amplified from N2 genomic DNA and Gibson cloned into a AvrII+SpeI double-digested pJW1592 vector (Ashley et al., 2020) as described in (Dickinson et al., 2015). N2 animals were injected and knock-ins were recovered as previously described (Dickinson et al., 2015)A previously described Cas9+sgRNA plasmid (pJW1254) (Zhang et al., 2015)targeting the 3’ end of *nhr-23* was used to generate the knock-in.

*nhr-23(wrd8[nhr-23::GFP^AID*::3xFLAG])* was generated with the self-excising cassette (SEC method (Dickinson et al., 2015) using a pJW1725 (nhr-23:GFP:SEC:degron:3XFLAG with *nhr-23* homology arms, U6p::sgRNA) repair template. The repair template (pJW1725) was constructed using SapTrap (Schwartz and Jorgensen, 2016) cloning with an SEC selection block (Dickinson et al., 2018). The 5’ homology arm was PCR amplified from a gBlock (IDT) and contained silent mutations to inactivate the PAM and two SapI restriction sites. The 3’ homology arm was PCR amplified from N2 genomic DNA. The 5’ and 3’ homology arm PCRs were cloned into pCR-Blunt II-TOPO using a Zero Blunt™ TOPO™ PCR Cloning Kit (Invitrogen, catalog #450245) to generate pJW1776 and pJW1781, respectively. pJW1725 was assembled through SapTrap cloning using pDD379 (backbone), pMLS287 (flexible linker), pDD372 (GFP), pDD363 (SEC), pJW1659 (linker::AID*::3xFLAG, pJW1776 (5’ homology arm), pJW1781 (3’ homology arm), and annealed oligos 3488+3489 (*nhr-23* targeting sgRNA) (Tables S1 and S2). The plasmid was injected into EG9615, which stably expresses Cas9, and knock-in animals were recovered as described (Dickinson et al., 2015). The strain was outcrossed two times to remove the *mex-5p::Cas9* allele and the SEC was excised to make JDW29.

*spe-44(wrd20[spe-44::30xlinker::mScarlet^3xMyc])* was generated using SapTrap cloning with an SEC cassette, similar to the approach described above. Oligos 3916+3917 were annealed and Sap Trap cloned (Schwartz and Jorgensen, 2016) into pJW1838 {Ashley:2020ib}to generate pJW1872(*U6p::sgRNA (F+E)* targeting *spe-44*). A pJW1876 repair template (30 amino acid flexible linker::mScarlet^SEC (Lox511I)^3xMyc with *spe-44* homology arms) was previously described (Ashley et al., 2020). The plasmid was injected into EG9615 and knock-in animals were recovered as described (Dickinson et al., 2015). The strain was outcrossed 2 times to remove the *mex-5p::Cas9* allele and the SEC was excised to make JDW101.

wrdSi10 was generated with the self-excising cassette method (Dickinson et al., 2015) and a recently described TIR::F2A::mTagBFP2 construct (Ashley et al., 2020). pJW1357 (*mex-5p::TIR1:F2A:mTagBFP2*) {Ashley:2020ib}and pCZGY2747 (*eft-3p::Cas9*; LGI ttTi4348 sgRNA vector) injected into JDW29 to make JDW39. *wrd10* was subsequently outcrossed to N2 animals one time to generate JDW84, then heat-shocked at 34 °C for 2-3 hours to excise the SEC to make JDW89. These two strains carrying a *mex-5p::TIR1:F2A:mTagBFP2* with and without the SEC, respectively are part of our recent AID toolkit and will be made available to the *Caenorhabditis* Genetics Center.

wrdSi12, 13 and 16 were made by crossing JDW39 to JK816 [fem-3(q20) IV], JK3107 {fbf-1(ok91) fbf-2(q704)/mIn1 [dpy-10(e128) mIs14] II}, and DR466 [him-5(e1490) V], respectively. *wrdSi19* was made by crossing KRY87 {nhr-23[kry61(nhr-23::AID*-TEV-3xFLAG)] I} to JDW83 [wrdSi10(mex-5p::TIR1:F2A:mTagBFP2:tbb-2 3’UTR+SEC, I:-5.32); him-5(e1490) V]. The Rol phenotype produced by expression of the sqt-1 gene within the SEC was used as a genetic marker to follow the insertion. Once the resultant strains were established, the SEC was excised by heat shock for 2-3 hours at 34 °C in those strains, thereby making JDW45, JDW47, JDW87 and JDW92, respectively.

#### Plasmid sequences are available upon request

JDW32 *[fog-3(wrd6)]* was made by ribonucleoprotein-based co-CRISPR injection into JDW29 (Arribere et al., 2014; Paix et al., 2015). sgRNAs were generated by *in vitro* transcription of PCR templates, as described by (Leonetti et al., 2016). JDW29 animals were injected with 3.8 uM dpy-10 sgRNA, 25.0 uM fog-3 sgRNA, 15.1 uM Cas9 protein, 0.4 uM dpy-10 repair oligo, 1.5 uM fog-3 repair oligo. The sequence of the sgRNAs and repair oligos is provided in Table S2. The insertion was targeted to the middle of exon 2 in the *fog*-3 locus and contained an EcoRI site to facilitate genotyping, followed by an in-frame stop codon. An additional base was inserted after the stop codon to produce a frameshift. F1 rollers were selected and screened by PCR and restriction digestion as described previously (Ward, 2015).

### Western Blot

Gravid JDW87 adults were allowed to lay eggs for 1 hour on MYOB with and without 4mM auxin (Indole 3-acetic acid, Alfa Aesar #AAA1055622), then removed. Progeny were allowed to grow until L4 + 1 day at 20°C. 100 hand-picked males were collected, washed 3X in M9 with 0.05% gelatin (VWR #97062-620) and resuspended in 30ul M9+gelatin. Worms were freeze/thawed 3X in liquid nitrogen and 10ul Laemmli sample buffer with 10% BME was mixed in. Samples were boiled for 10 minutes at 95 °C, centrifuged for 5 minutes at full speed and the supernatant was removed to a new tube. Proteins were separated on a SDS-PAGE Mini-PROTEAN TGX 4-15% gradient gel (Bio-Rad, catalog #456-1086) along a 1:1 mix of Amersham ECL Rainbow Molecular Weight Markers (Amersham # 95040-114) and Precision Plus Protein™ Unstained Standards #1610363. Proteins were transferred to a PVDF membrane using a Trans-Blot Turbo Transfer System. Membrane was washed in TBST and blocked in TBST + 5% milk (VWR # 10128-600) for 1 hour at room temperature. Membrane was incubated in TBST + 1:1000 Anti-FLAG-M2-HRP (Sigma, #8592) at 4°C overnight while rolling, washed 5X in TBST and incubated in 10ml Supersignal West Femto Maximum Sensitivity Substrate (Thermo Scientific Pierce #34095) for 5 minutes. Separated proteins on the gel, transferred proteins on the PVDF and the final blot were imaged using the “chemi high-resolution” setting on a Bio-Rad ChemiDoc MP System.

### Auxin Depletion

Control media consisted of MYOB + 0.25% ethanol. Indole 3-acetic acid was dissolved in 100% ethanol to 1.6M and mixed into MYOB to 4mM to make auxin media. For live progeny counts, gravid adults were allowed to lay F1 eggs for 1 hour on control or auxin media and then removed. F1 progeny grew at 20°C until L4, then were singled onto equivalent media to lay eggs for 24 hours (n=12 per condition). These worms were transferred to a new and equivalent set of media each day for 3 days and F2 eggs were counted. Once the F2 progeny of these worms grew to L4, they were counted. For matings, F1 progeny were grown to L4, then singled or mated with L4 males for 24 hours at 20°C. Transfers and counts were then performed as described. To determine the auxin-sensitive period of development, F1 progeny eggs were laid on MYOB media + ethanol with or without 4mM auxin, allowed to grow until the appropriate stage and were transferred to the opposite plate-type for the duration of their development. Once F1 progeny reached L4 + 1 day, transfers began as described and F2 progeny were counted. For cell morphology and immunohistochemistry, worms were grown from the late L1 stage on 4mM auxin media.

### Co-suppression

*nhr-23* cDNA was PCR amplified and Gateway cloned (Invitrogen) into pDONR221 to produce pJW258. The *mex-5* promoter was amplified from *C. elegans* N2 genomic DNA and an *nhr-23* cDNA was amplified from pJW258. PCR stitching was used to create a chimeric *mex-5p::nhr-23* cDNA PCR product, as previously described (Dernburg et al., 2000). 50 ng/μl was injected into JDW45 *[fem-3(q20)]* hermaphrodites along with 10 ng/ul pCFJ90 *[Pmyo-2::mCherry::unc-54utr]*, a pharyngeal-expressed co-injection marker. Worms were incubated at 25°C to masculinize the germlines of the subsequent F1 progeny. *myo-2p::mCherry* positive F1 young adults were then imaged for defective/arrested spermatocytes.

### Statistical analyses

Statistical tests and numbers of animals analyzed are detailed in figure legends.

## Acknowledgements

This work was funded by the National Institute of Health (NIH) National Institute of General Medical Sciences (NIGMS) [R00GM-107345 to JDW and R15GM-096309 to DCS]. Some strains were provided by the *Caenorhabditis* Genetics Center, which is funded by the NIH Office of Research Infrastructure Programs [P40 OD010440]. We thank Diana Chu and Penny Sadler for critical reading of this manuscript.

## Author Contributions

J.M.R., R.M.M., D.C.S., and J.D.W. designed the experiments. All authors performed the experiments. J.M.R., A.L.A, D.C.S., and J.D.W. performed the data analyses. J.M.R., K.N.M., D.C.S., and J.D.W. wrote the paper with feedback from the other authors.

---

**Fig. S1. NHR-23-depletion in worms expressing TIR1 from a *sun-1* promoter causes sterility similar to that observed when TIR1 is expressed from the *mex-5* promoter.** (A) Count of L1 larvae and fertilized eggs or unfertilized oocyte progeny from parents grown from L1 onwards with or without 4mM auxin. Error bars=standard deviation (n=12 for each condition) (B) Representative DIC images of L4 + 1 day *nhr-23::GFP::Biotag::AID*::3xFLAG; sun-1p::TIR1* hermaphrodites.

